# Protein flexibility and dissociation pathway differentiation can explain onset of resistance mutations in kinases

**DOI:** 10.1101/2021.07.02.450932

**Authors:** Mrinal Shekhar, Zachary Smith, Markus Seeliger, Pratyush Tiwary

**Affiliations:** Center for Development of Therapeutics, Broad Institute of MIT and Harvard, Cambridge, MA, USA; Biophysics Program and Institute for Physical Science and Technology, University of Maryland, College Park 20742, USA; Department of Pharmacological Sciences, Stony Brook University, Stony Brook, New York 11794-8651, USA; Department of Chemistry and Biochemistry and Institute for Physical Science and Technology, University of Maryland, College Park 20742, USA

## Abstract

Understanding how point mutations can render a ligand or a drug ineffective against a given biological target is a problem of immense fundamental and practical relevance. Often the efficacy of such resistance mutations can be explained purely on a thermodynamic basis wherein the mutated system displays a reduced binding affinity for the ligand. However, the more perplexing and harder to explain situation is when two protein sequences have the same binding affinity for a drug. In this work, we demonstrate how all-atom molecular dynamics simulations, specifically using recent developments grounded in statistical mechanics and information theory, can provide a detailed mechanistic rationale for such variances. We establish the dissociation mechanism for the popular anti-cancer drug Imatinib (Gleevec) against wild-type and N387S mutant of Abl kinase. We show how this single point mutation triggers a non-local response in the protein’s flexibility and eventually leads to pathway differentiation during dissociation. This pathway differentiation explains why Gleevec has a long residence time in the wild-type Abl, but for the mutant, by opening up a backdoor pathway for ligand exit, an order of magnitude shorter residence time is obtained. We thus believe that this work marks an efficient and scalable approach to pinpoint the molecular determinants of resistance mutations in biomolecular receptors of pharmacological relevance that are hard to explain using a simple structural perspective and require mechanistic and kinetic insights.

**Significance statement:** Relapse in late-stage cancer patients is often correlated with the onset of drug resistance mutations. Some of these mutations are very far from the binding site and thus hard to explain from a purely structural perspective. Here we employ all-atom molecular dynamics simulations aided by ideas from information theory that can reach timescales of seconds with minimal human bias in how the sampling is enhanced. Through these we explain how a single point mutation triggers a non-local response in the protein kinase’s flexibility and eventually leads to pathway differentiation during dissociation, thereby significantly reducing the residence time of the drug.

## Introduction

Post-translational modifications of protein kinases play a crucial role in the modulation of multiple signaling pathways, which regulate key cellular processes like cell growth and proliferation. ^1,2^ Protein kinases assist phosphorylation by catalyzing the transfer of *γ*-phosphate of an ATP molecule to the hydroxyl group of Ser, Thr, or Tyr residues.^1,2^ Kinases thereby act as switches that control key cellular signaling pathways. Over the years, the importance of kinases as drug targets and thereby effort to develop therapeutic agents for modulating kinase activity has been well established. ^2–4^ The development of kinase inhibitors is however challenging because of the high sequence conservation of the kinase ATP-binding site, the major site targeted by these small molecules. However, despite the challenges, remarkable progress in the field of kinase-based drug design was made with the discovery of Imatinib (Gleevec)^5,6^ as a potent Abl kinase inhibitor. Gleevec has continually been found to be highly efficacious in the treatment of early-stage chronic myeloid leukemia (CML). ^5,6^ Unfortunately, a large fraction of late-stage cancer patients suffer from cancer relapse due to the onset of drug resistance.^7–10^

Understanding the molecular basis of the effects of these oncogenic mutations on the efficacy of cancer drugs is the first step in solving the problem of drug resistance. From a molecular point of view, the mutations can be classified as either orthosteric i.e, in the inhibitor binding site directly affecting Imatinib binding or allosteric wherein these mutations modulate drug resistance indirectly.^11^ In the past, multiple computational^7,12,13^ and experimental^11,14^ studies have tried to explain the resistance mechanism by either pointing to direct abrogation of H-bonding interactions or steric effects in the reduction of the binding affinity. The well-studied T315I mutation in Abl kinase is such an example. ^12,15,16^ Particularly, alchemical free energy-based calculations ^17,18^ have been employed to quantify the direct effects of the point mutation on inhibitor binding free energy. Although alchemical methods to a certain degree have been able to quantify the effects of resistant mutations on inhibitor binding free energies, it remains untenable to use them for allosteric mutations that rely mainly on modulating conformational dynamics of the kinase. Of particular interest are a certain class of mutations that have no appreciable effect on Imatinib binding affinity. Rather, these mutations are expected to regulate drug efficacy by modulating drug dissociation kinetics. Thus, for this class of mutations understanding the drug dissociation kinetics is paramount to understanding drug resistance. Despite the importance of kinetic measurements in understanding drug efficacy,^19–21^ such measurements either computationally or experimentally have remained non-trivial. In the context of drug-receptor binding, experimental methods have been useful in the elucidation of low energy states, however, they have lacked in the description of short-lived metastable and transition states.

Our specific interest in this work pertains to a recent discovery made by Lyczek etitat al,^22^ who have identified a novel N387S mutant of Abl kinase to which the FDA-approved and widely prescribed oral chemotherapy drug Imatinib (Gleevec) binds with a similar binding affinity as it does to the wild-type (WT). However, they find that Imatinib displays a three-time faster dissociation rate *k_off_* from the mutant than from the WT. In this work, we use recent statistical mechanics and information theory-based all-atom resolution methods^23–25^ to provide a mechanistic all-atom resolution rationale for this perplexing and important finding. Our method provides absolute *k_off_* estimates for both WT and mutant Abl within an order of magnitude of the experimentally determined values, also capturing their relative magnitudes. Going even beyond reproducing *k_off_* values, our calculations directly pinpoint the varied dissociation pathways and mechanisms adopted by Imatinib in both variants of Abl kinase. Our key mechanistic finding can be summarized as follows: We find that there are two distinct Imatinib dissociation pathways for the WT and the mutant Abl. Furthermore, we explain the order of magnitude difference between WT and mutant Imatinib unbinding kinetics by invoking varied structural rearrangements required to allow for two distinct Imatinib dissociation pathways in WT and mutant Abl.

To perform such mechanistically insightful simulations, in principle one could use computational methods like molecular dynamics (MD) simulations. These have the potential to not just quantify k_*off*_, but also give a direct atomistic understanding of metastable states along with drug binding/unbinding pathways which have remained elusive through experimental measurements. However, even with highly specialized hardware, brute-force MD has been limited to at best a fraction of a millisecond.^26,27^ On the other hand, numerous enhanced sampling algorithms ^28–38^ have been employed for k_*off*_ calculations with more reasonable computational costs and the ability to achieve pharmacologically relevant timescales of seconds, minutes, and slower. Here we use one such enhanced sampling method “infrequent metadynamics” that has been employed to obtain unbiased estimates of dissociation kinetics in numerous systems.^23,35,36,39^ The reliability of such calculations ^40^ is closely linked to the ability to design an appropriate reaction coordinate (RC) that describes the dissociation process and that can be used to construct the low-dimensional biasing potential as a function thereof. In general, learning such an RC is an extremely difficult problem especially for rare events such as drug dissociation, wherein any framework used to learn the RC depends on the quality of the sampling, while accurate sampling itself can not be achieved without having a reasonable RC. In this work, we make use of recently developed machine learning and statistical mechanics-based methods^23–25^ that tackle the above chicken-versus-egg problem through systematically iterating between sampling and RC optimization in a nearly automated manner (see Fig. 6 in Supplementary Information (SI) for a flowchart of the overall protocol). Once a reliable RC is obtained, infrequent metadynamics can be performed along it to calculate k_*off*_ estimates with error bars ^41^ and directly observe the entire dissociation pathway with all-atom and femtosecond resolution.

## Results and discussion

### Abl-kinase structure

For the sake of completeness, we begin by summarizing some well-known structural details. Abl kinase has a typical bilobal kinase domain, consisting of a smaller N-terminal lobe (N-lobe) and a larger C-terminal lobe (C-lobe). The ATP binding site is located between the N-lobe and the C-lobe of the catalytic domain, distinguished by conserved structural features like the phosphate-positioning loop (P-loop), the activation loop (A-loop), the *α*C helix, and the conserved Asp-Phe-Gly (DFG) motif (Fig. 1). As evidenced by biophysical experiments^42–45^in addition to computational ^46–53^ and structural studies, ^8,12,44,54–56^ Abl-kinase exists in a dynamic equilibrium between multiple conformations: active, inactive and multiple intermediate states interspersing these. These states are characterized by the conformational flexibility of evolutionarily preserved kinase motifs such as the A-loop, DFG motif and the *α*C-helix (Fig. 1). Abl-kinase in its active conformation is characterized by the “DFG-in” and “*α*C-in” state. In this state the D400 residue of the DFG loop points towards the active site, ready to coordinate with ATP and the Mg^2+^ ion. The *α*C helix swings in towards the binding site adopting the “*α*C-in” conformation and allowing for ion-pair interaction between the conserved E305 on *α*C helix with conserved K290 in the *β*3 strand. At the same time, the A-loop adopts an extended conformation allowing for space for the substrate to dock.

**Figure 1:**
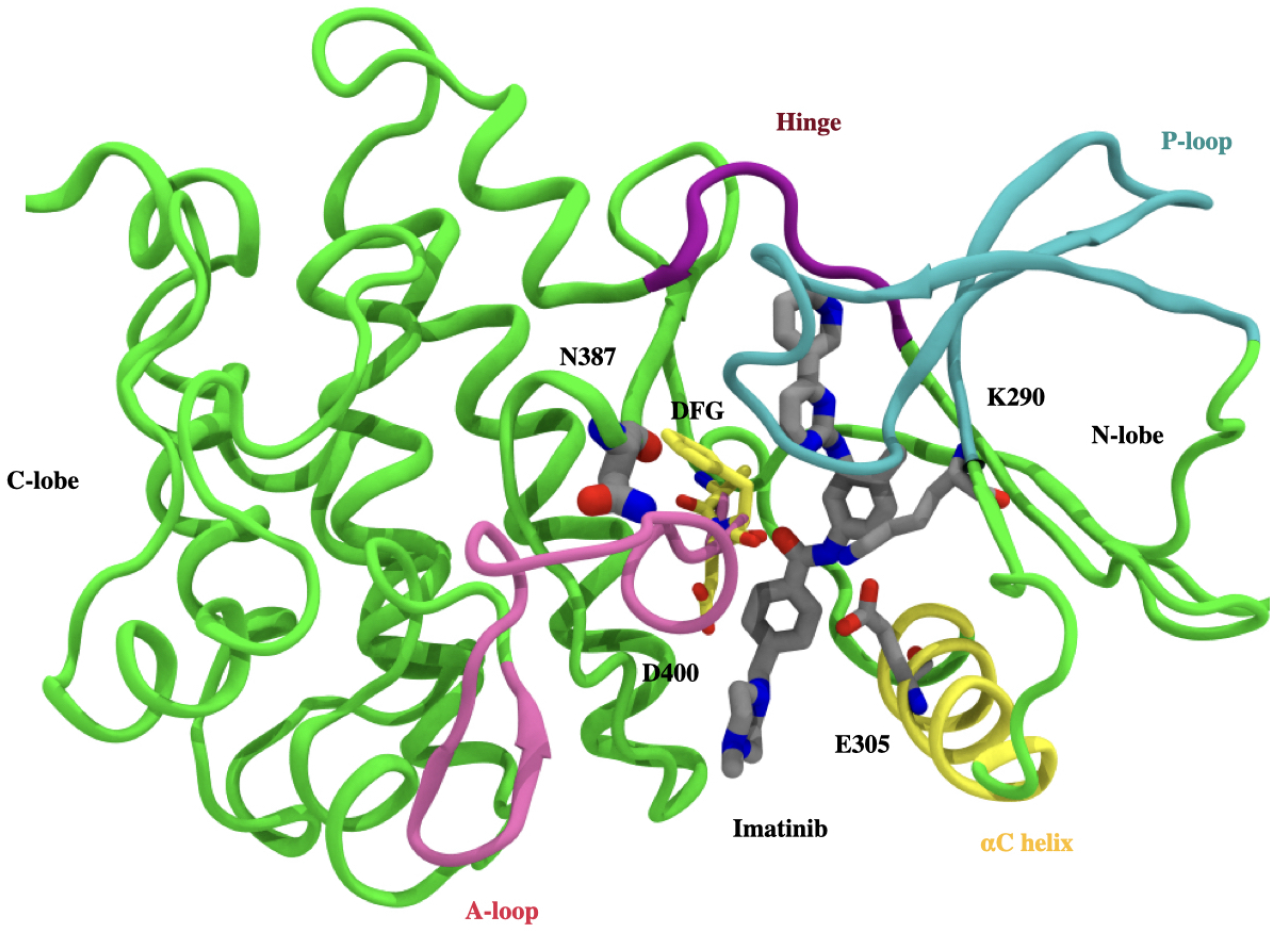
Crystallographic binding mode: Imatinib (grey) bound to the catalytic domain of Abl kinase (PDB id 1OPJ). Kinase domain is divided into N-terminal lobe (N-lobe) and C-terminal lobe (C-lobe), with the inhibitor (Imatinib) binding site located between the lobes. Abl-kinase has conserved structural features like A-loop (pink ribbon), P-loop (blue ribbon), *α*C helix (yellow ribbon), hinge (purple ribbon), and DFG motif (yellow sticks). Salt bridges involve D305 in the DFG motif, R386 in the A-loop, E305 in the *α*C helix, and K290 cover and block the binding tunnel in front. The mutant residue N387S lies behind the DFG motif and is shown in a gray stick.

The conserved E305, in turn, interacts with the *α* and *β* phosphates of ATP. In contrast to this very specific characterized active state, a kinase can also adopt one of many inactive conformations. ^57–60^ One such often-mentioned conformation is the so-called “DFG-out” conformation, where the D400 switches its position with the phenylalanine F402, pointing away from the ATP binding site. Type II inhibitors such as Imatinib are suspected to selectively target this “DFG-out” inactive state.^14,61^ Imatinib is stabilized in the ATP binding site predominately via hydrophobic interactions along with 6 hydrogen bonding interactions with the side chain hydroxyl of the gatekeeper residue T315, backbone and side chain of E305, and the carbonyl and backbone-NH of D381 on the *α*C helix.

### Infrequent metadynamics simulations reveal two distinct pathways

In this work, our central aim is to use all-atom MD simulations to understand the molecular determinants that result in an order of magnitude difference in the measured Imatinib k_*off*_ for WT relative to the mutant N387S Abl kinase. Due to the extremely slow timescales of the dissociation process relative to what can be achieved in MD simulations, here we use a combination of enhanced sampling methods as mentioned in the Introduction and detailed in the Methods section. The first step in the k_*off*_ calculation for Imatinib is the description of a low-dimensional reaction coordinate (RC) that can suitably describe biological processes of interest, in this case, the dissociation of Imatinib from WT/Mutant Abl kinase. Briefly, we perform 100 ns long unbiased MD simulation on Imatinib bound with WT and Mutant Abl kinase respectively. By employing the method AMINO^25^ (Methods) on these trajectories we learned non-redundant trial order parameters (OPs) that describe Imatinib dissociation (Fig. 2 A-B). Starting from a total possible 84 OPs corresponding to different possible kinase-Imatinib heavy-atom contacts, AMINO identifies 5 OPs to be sufficient for the WT and the mutant complexes respectively shown in Fig. 2 A-B. See Table 2 in SI for a summary of OPs. These OPs are expressed as distances between C*α* atoms of highlighted residues and centers of mass of two halves of Imatinib. These two halves are labeled p2a (blue stick), which is initially solvent exposed, and p2b (orange stick) which is the buried half. In the case of WT, AMINO derived OPs measure the distance between p2a and the C*α* atoms of the residues Y251, Y272 along with the distance between p2b and C*α* atoms of the residues E257 and S248. In comparison, mutant OPs are defined by the distance between p2a and the C*α* atoms of the residues I332,K293 along with the distance between p2b and C*α* atoms of the residues D344,Y272, F302. Along with these OPs, the other common OPs for mutant and WT are the distances between C*α*C*β* atoms of T334 shown in red spheres and the N3 and N4 atoms of Imatinib shown in blue spheres. Subsequently, multiple rounds of maximum caliber-based SGOOP optimization^24,62^ are performed to construct an even lower-dimensional RC from these OPs shown in Fig. 2 C-D. Finally, 11 independent trials of infrequent metadynamics were performed starting from the bound crystal pose and biasing the aforementioned RC, stopping when the ligand was fully solvated. Through these, we generated an ensemble of ligand dissociation pathways along with the associated protein conformational changes and culminating in the k_*off*_ calculation (Fig. 2 E) through the procedure of Ref. ^23,39^

**Figure 2:**
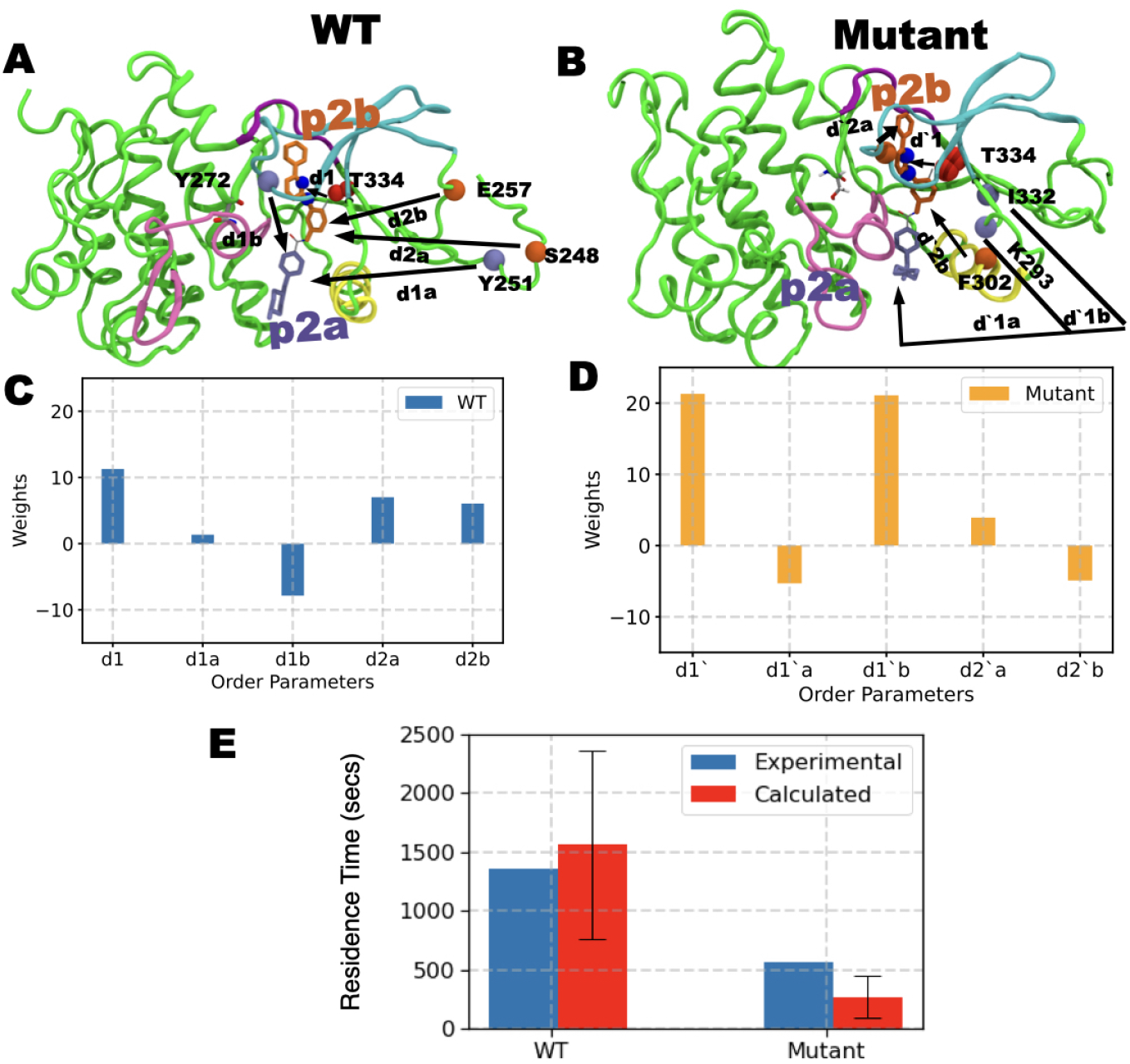
A) and B) show order parameters OPs obtained through AMINO for WT and Mutant respectively. C) and D) show the respective RC constructed from these OPs through SGOOP. E) shows the residence times (red bar) obtained by biasing the respective RC for WT and mutant system compared against experimental measurements (blue bars). The red bars represent the fitted residence time^39^ while the standard deviation is shown as a black error bar. See main text for further details of the OPs.

In the infrequent metadynamics simulations, depending upon the protein, WT or mutant, two distinct Imatinib release pathways were observed. The two pathways as shown in Fig. 3A (left/right upper panel), can be described as: (i) exit under the P-loop closer to the *α*C helix (i.e., through the hydrophobic pocket), henceforth named as the *α*C pathway, or (ii) exit through a pathway closer to the P-loop and the kinase hinge, named as the hinge pathway. These two pathways can be more quantitatively characterized by projecting them onto a 2-D space spanned by the two distances between the Imatinib center of mass and the centers of mass (i) of the alpha carbons of the hinge, and (ii) of the *α*C helix. Evidently, as can be seen in Fig. 3 A (upper/bottom left panel), in the WT dissociation trajectory, the hinge pathway is the dominant pathway. Quantitatively, Imatinib during its dissociation comes as close as 3.8±1 Å to the hinge region while not getting closer than 9±0.5 Å from the *α*C helix. Sharply contrasted to this, for dissociation from N387S we find that Imatinib does not get closer than 8±0.5 Å to the hinge region while it comes as close as 6±0.5 Å to the *α*C helix. These distances are the averaged values over the different independent trajectories with error bars shown in Fig. 3.

**Figure 3:**
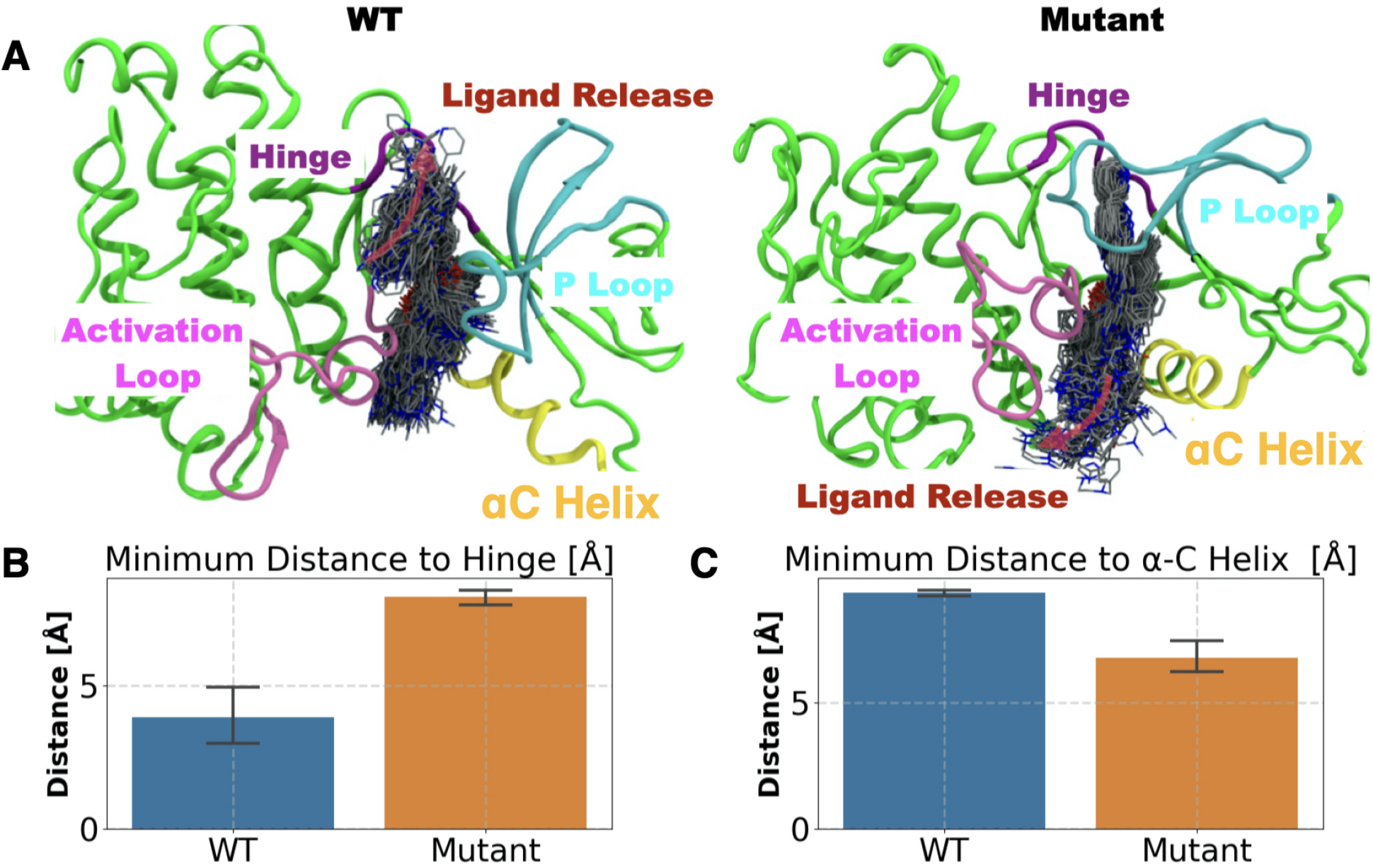
Mutational effect on the substrate release pathway (red arrows in the top panel): A) WT (top left) and Mutant (top right) panels. Ligand dissociation trajectory is depicted by ligands in licorice representation sampled every 100 ps. The overall direction of the substrate release is depicted by transparent pink arrows. Bar plot and the error bar representing the mean and standard deviation of the closest distance between the center of mass of Imatinib and B) hinge and C) *α*C helix for WT (blue bar) and mutant (orange bar) trajectories.

Together these observations give mechanistic insight into the different dissociation time scales for Imatinib from WT Abl and from N387S Abl. A single point mutation results in divergent pathways for Imatinib dissociation, opening up the possibility to take a much quicker release route for the drug and thus lowering its residence time. In the next section, we provide calculated residence times from our simulations for both systems along with experimental benchmarks, following which we provide an atomistic underpinning for the observed differences in the WT and mutant dissociation pathways and k_*off*_ values.

### Overall kinetics of the dissociation process

In order to determine the kinetics of the dissociation process, we first define the dissociated state of the ligand as when Imatinib has reached the solvent-exposed surface of the protein. We find that starting here there could be other “trap states” on the surface where Imatinib could bind with much weaker strength, but we ignore the kinetics corresponding to these as they can be expected to contribute much less relative to the time Imatinib takes to dissociate from the main binding site. More quantitatively, Imatinib is considered to be dissociated if the distance between the ligand center of mass and the binding site exceeds 15 Å. As stated in the previous section, ligand dissociation was observed in each of the 11 infrequent metadynamics simulations for the WT and for the mutant. By fitting the respective 11 observations of the residence time to a Poisson distribution as per the protocol in Ref.^36,39^ and described further in Supplementary Information (SI), we calculate the residence times *τ* for both systems along with 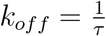. We also perform a p-value analysis (Table 1 and Fig. 8 in SI) for both which indicates that the residence time calculations meet the reliability threshold from Ref.^36,39^ The residence time for Imatinib in WT Abl was 1560±797 sec, in excellent agreement with the experimentally determined residence time of 1357 sec. On the other hand, we find a shorter residence time for Imatinib in the mutant Abl kinase, equalling 269±176 sec and again in excellent agreement with the experimentally measured value of 562 sec. Thus as can be seen in Fig. 2 E, qualitatively and quantitatively we are successful in recapitulating the experimentally observed residence times, with Imatinib k_*off*_ for mutant Abl being around an order of magnitude faster than WT. More importantly, apart from having a qualitative agreement in predicting the order of magnitude difference in Imatinib k_*off*_ for mutant vs WT Abl, the absolute values of the calculated residence time and k_*off*_ are well within the same order of magnitude as the experimentally determined values (Fig. 2 E). In the next section, the different dissociation pathways are explored in detail, describing the structural features distinguishing the two main pathways and finally culminating in providing a mechanistic explanation for the observed difference in Imatinib k_*off*_ values between WT and N387S Abl kinase.

**Table 1:** Summary of Imatinib dissociation kinetics, comparing the residence time/k_*off*_ of WT and mutant Abl as measured in this work against experiments in Ref.^22^

| System | Infreq<br>Metad<br>residence time (sec) | Infreq.<br>Metad.<br>$k_{off}(\text{sec}^{-1})$ | Experimental<br>residence time (sec) | Experimental<br>$k_{off}(\text{sec}^{-1})$ | p-value |
| --- | --- | --- | --- | --- | --- |
| Wild-Type | $1560 \pm 797$ | $6.41 \pm 3.28 \text{ e-04}$ | 1357 | $7.368 \text{ e-04}$ | 0.11 |
| N368S | $269 \pm 176$ | $3.700 \pm 0.24 \text{ e-03}$ | 562 | $1.770 \text{ e-03}$ | 0.23. |

### Mechanistic underpinning of mutational effects on dissociation path

In the previous sections, we established that Imatinib unbinds from WT-Abl and N387S-Abl dominantly via the hinge pathway and the *α*C pathway respectively and that our enhanced sampling approach is based on AMINO, SGOOP and infrequent metadynamics can return quantitatively accurate residence times for Imatinib in WT and N387S Abl kinase. We now provide further mechanistic analysis joining the results of the previous sections. For this, we approximately classify the Imatinib-Abl kinase system in three states, on the basis of the distance *d* between the Imatinib center of mass and the binding site. These are the (i) starting state (crystal structure), (ii) pre-release state (*d* ≤ 4.5 within Å) and (iii) the dissociated state (d≥15 Å).

We observe that for Imatinib in complex with the WT-Abl system (Fig. 4 A), the substrate release pathway is enclosed by a triad of interactions between the P-loop, hinge, and the DFG loop. In this state, the residue Y272 from the P-loop forms a hydrogen bond with N341 from the hinge. Furthermore, Y272 packs against F401 from the DFG loop stabilized by the CH–*π* interaction. Subsequently, as Abl kinase transits to the pre-release state, P-loop moves away from the hinge and the DFG loop, resulting in the loss of the Y272-N341 hydrogen bond. The resultant pre-release state is characterized by weakened P-loop hinge interaction and an open pathway for substrate release. Consistent with this, for the WT Abl-Imatinib system by averaging over all 11 independent dissociation trajectories, we find (Fig. 4 A, C) the minimum attained distance between Imatinib and the hinge region to be 8.7±3.1 Å, implying an open pathway for the substrate release proximal to the P-loop-hinge region. In comparison to the dissociation of Imatinib from N387S Abl, the corresponding Imatinib-Hinge region minimum distance (Fig. 4 C) is 6±3.73 Å, signifying a relatively occluded P-loop-hinge pathway as compared to WT Abl. In the case of N387S mutant Abl, *α*C pathway is blocked by the packing of *α*C helix against the DFG motif and Imatinib. Particularly, the residue K290 from the K290–E305 of the prototypical ionic lock interacts with the backbone carbonyl (Fig. 4 B) of the residue D400 from the DFG motif. Furthermore, the backbone N-H of the F401 from the DFG motif stabilizes the binding of Imatinib. As mutant Abl transits to the pre-released state, the major conformational change that we observe involves (Fig. 4 B) the outward motion of *α*C helix from the *α*C helix-in state. As a direct consequence of the *α*C helix outward motion, we observe diminished K290-DFG interaction resulting in the formation of an open the *α*C helix pathway. Consistently, in the N387S Abl-Imatinib system by averaging over all 11 independent dissociation trajectories, we find (Fig. 4 D) the K290-D400 distance to be 7.9±1 Å in the pre-released state. Comparatively, for WT Abl-Imatinib, the average K290-D400 distance was 5±3 Å, signifying an open *α*C helix pathway for Imatinib release.

**Figure 4:**
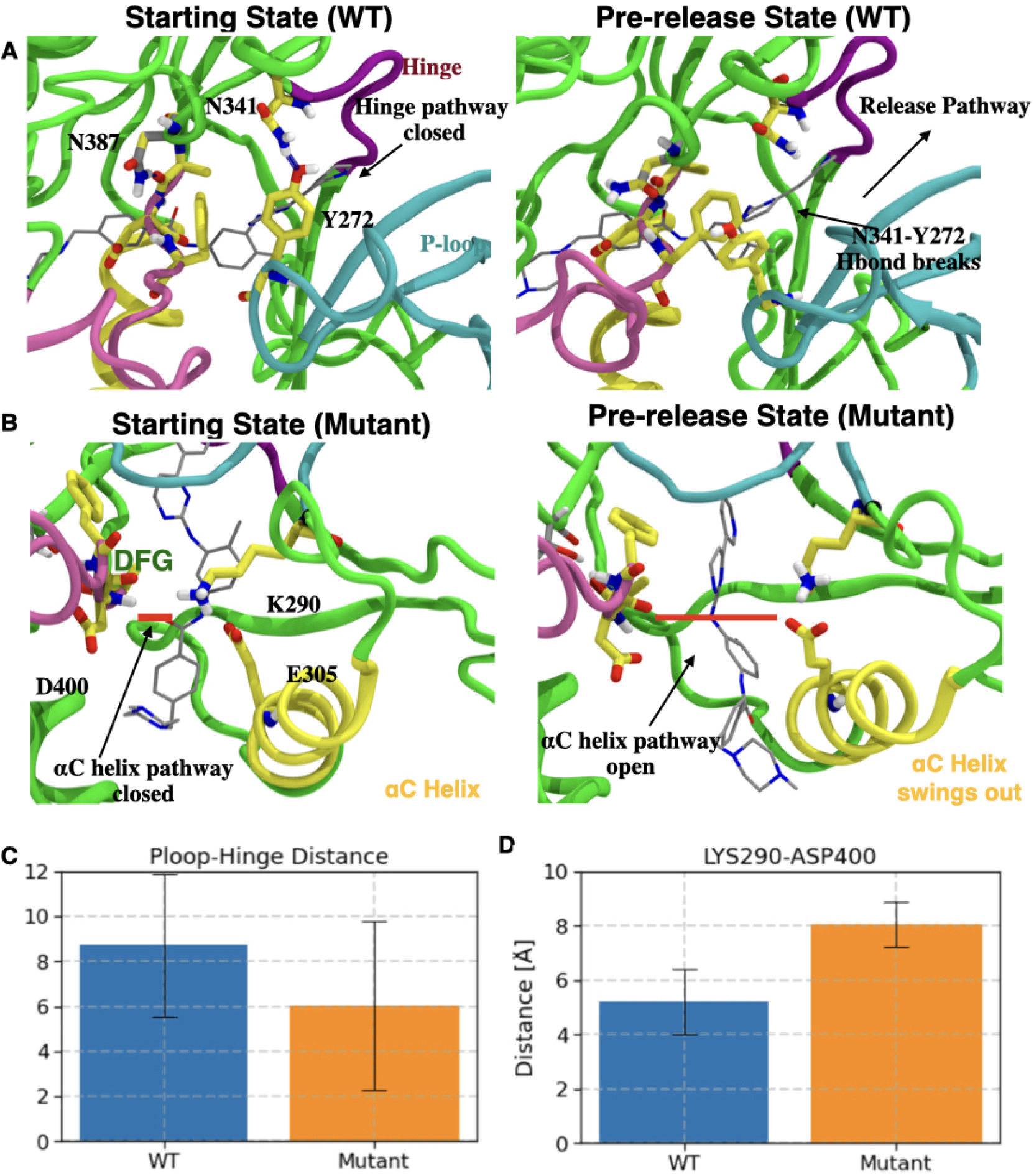
A) Left: Starting state of WT Abl (Crystal structure). H-bond between Y272 (P-loop) and N341 (hinge) is shown as a blue dashed line. Right: Abl conformation transiting to the pre-release state involves disrupted h-bond between Y272 and N341 forming an open ligand release pathway depicted by a black arrow. B) Left: H-bond between D400 from the DFG motif and Imatinib (gray sticks) in N387S Abl is shown as a thick red line. Right: Conformational changes in N387S Abl as it transits to the pre-released state involve the disruption of the interaction network of E305, K290, Imatinib, and DFG motif. C) and D) respectively show the closest distance (with error bars) between C) C*α* atoms of Y272 and N341 for WT Abl (Blue) and N387S Abl (orange); and D) C*α* atoms of K290 and D400 for WT Abl (Blue) and N387S Abl (orange).

In order to further understand the molecular basis of distinct Imatinib release pathways for WT and N387S Abl and the resulting differences in dissociation kinetics, we also analyze the effect of mutation on the conformational dynamics of the DFG motif. We observe (Fig. 5 A (left panel)) that the residue N387 in WT Abl forms hydrogen bonds with the backbone of residue A399 which is one residue upstream of D400 from the DFG motif. However, in N387S Abl, the mutated residue Serine with a smaller side-chain than the original Asparagine in the WT Abl has diminished propensity (Fig. 5 A (right panel)) to form the aforementioned hydrogen bonding interaction. Evidently, by averaging over all independent 11 WT dissociation trajectories prior to the substrate release, we observe (Fig. 5 B) that on average N387 forms 0.8±0.5 hydrogen bonds with backbone A399. On the other hand, by averaging over all independent 11 trajectories, the average number of hydrogen bonds between S387 and DFG motif was observed to be only 0.3±0.5. As a direct consequence of impaired hydrogen bonding interactions, we observe that the DFG motif has an elevated conformational flexibility (Fig. 5 B) in mutant Abl as compared to WT. Consistently, we observe that on an average in the N387S Abl-Imatinib trajectories, the mean root mean square deviation (RMSD) of the DFG motif from the crystal structure was observed to be 2.8 Å in comparison to 2.2 Å for the WT trajectories.

**Figure 5:**
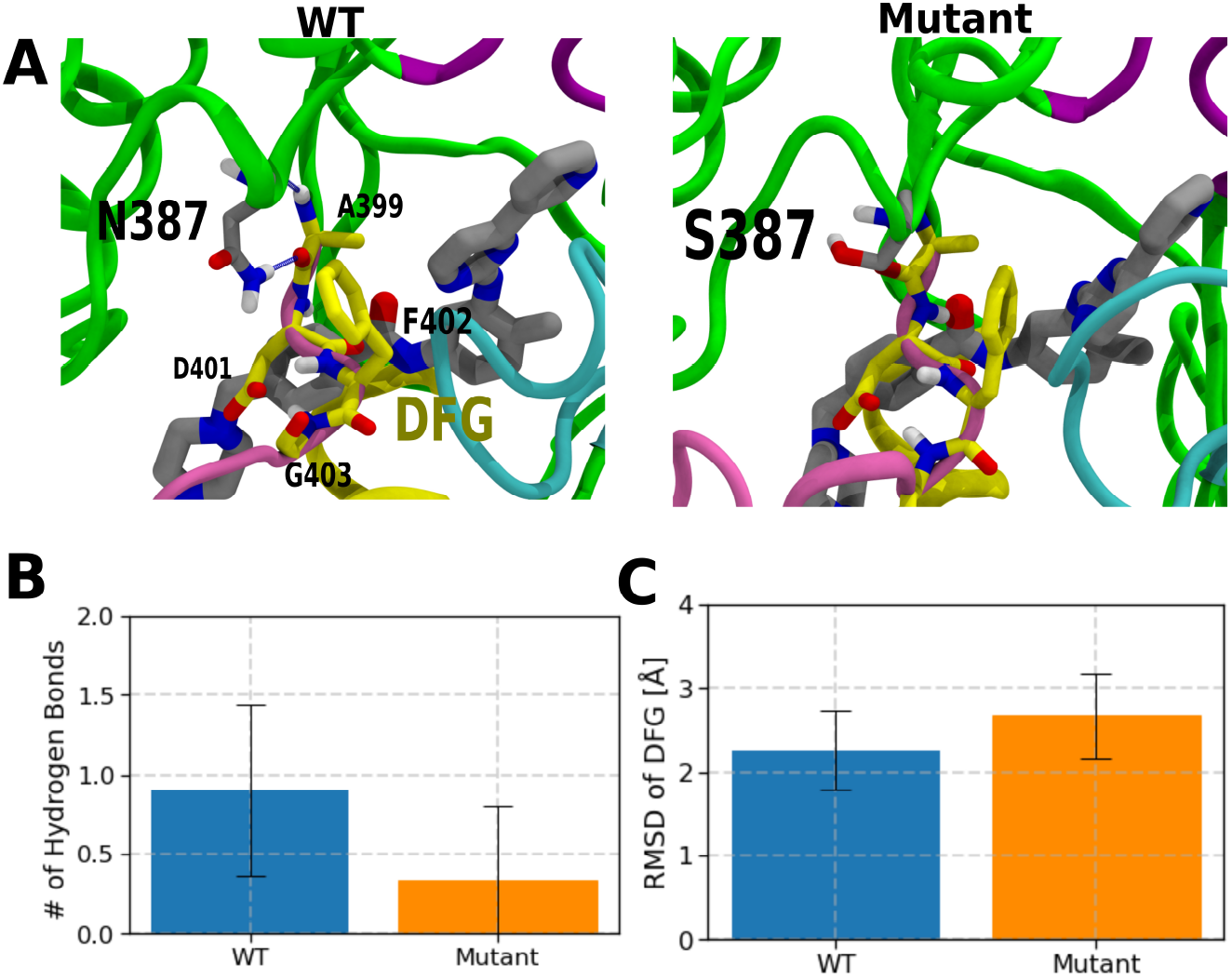
Flexibility of DFG motif is modulated by h-bond interaction of N387 and the DFG loop. A) Crystal structure pose of Imatinib (gray licorice representation) in WT (left panel) and N387S (right panel). DFG motif is represented by yellow licorice. H-bond between N387 (gray licorice) and A399 is shown as blue dashed line. Bar plot and the error bar representing the mean and standard deviation of B) N387 and DFG motif for WT Abl (Blue bar) and N387S Abl (orange bar) C) Root mean square deviation (RMSD) of DFG motif from the crystal structure position for WT Abl (Blue bar) and N387S Abl (orange bar).

**Figure 6:**
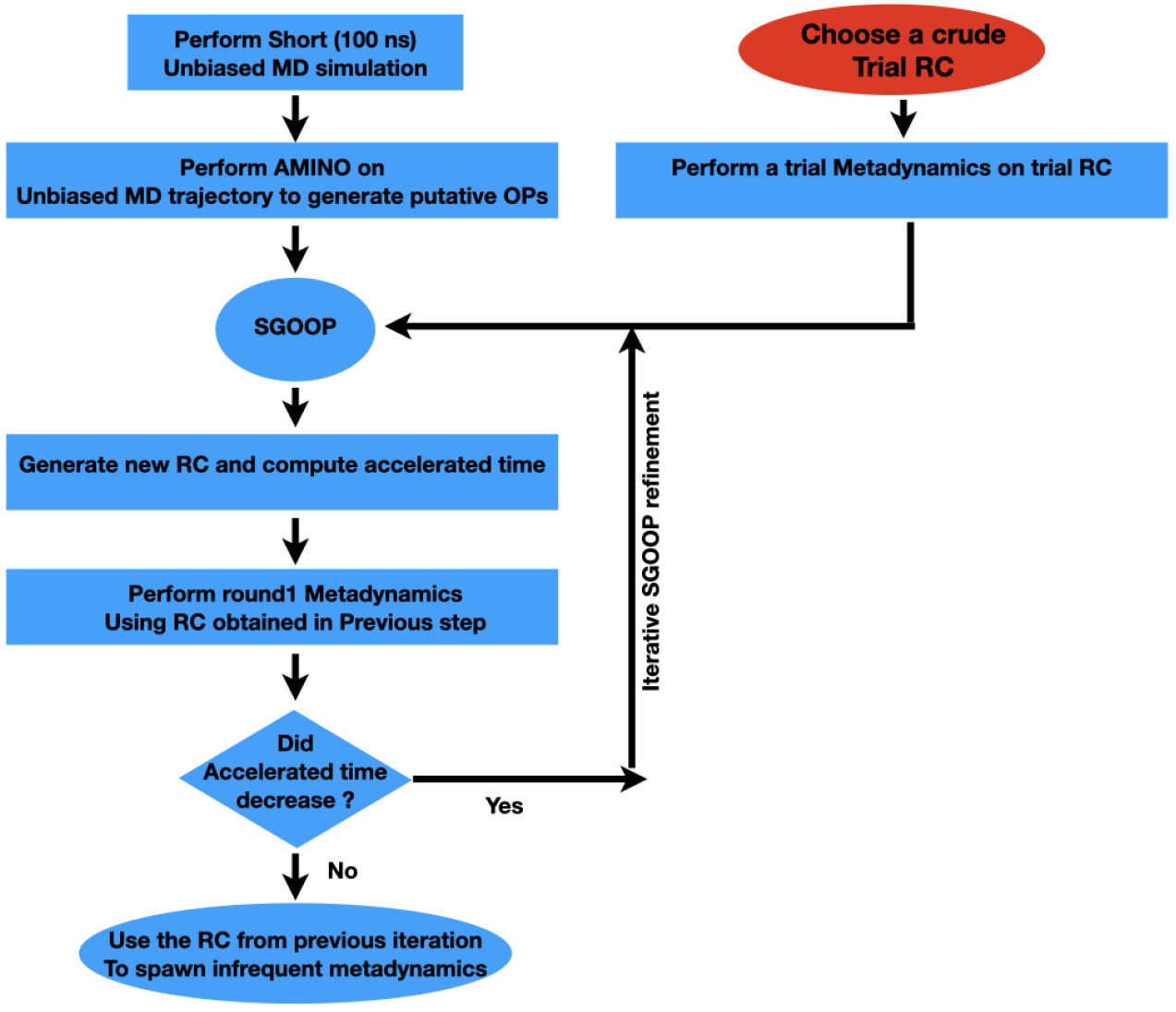
Schematic representing our protocol to generate a suitable RC to be used in biasing infrequent metadynamics simulation. Starting from a trial RC and an unbiased MD simulation, we perform AMINO to generate a list of OPs. Subsequently, we systematically iterate between sampling and RC optimization in a nearly automated manner to obtain the RC to used to bias infrequent metadynamics simulations.

We hypothesize that the increased flexibility of the DFG motif in the mutated N387S Abl kinase correlates with the diminished interaction between K290 and the DFG backbone, thereby allowing for the outward *α*C helix motion. This outward *α*C helix motion results in an open Imatinib release pathway proximal to *α*C helix. However, the P-loop and hinge interaction in mutant Abl remains unaffected and hence the pathway proximal to the hinge remains closed. In comparison to WT Abl, the DFG motif has diminished conformational flexibility, and thus the K290 DFG interaction is conserved. As a result, the *α*C helix remains packed against the DFG motif effectively blocking the *α*C helix proximal pathway. At the same time, we observe that in WT Abl instead of the outward *α*C helix motion, the P-loop and hinge interaction is weakened by the outward motion of the P-loop from the binding site. The outward P-loop motion contributes to opening up the substrate release pathway proximal to the hinge rather than the *α*C helix.

## Conclusion

Drug resistance in late-stage cancer patients remains a major factor in the failure of anti-cancer therapeutic treatments. In this work, we describe an efficient formalism to characterize the molecular determinants of the resistance mutations, leading to potentially develop therapeutics that can potentially overcome the drug efficacy loss due to resistance mutations. Particularly, we focus on a novel N387S Abl kinase mutation discovered by Lyczek *et al* ^22^ that results in a three times faster k_*off*_ for Imatinib against mutated Abl versus wild-type Abl, while the Imatinib binding affinity remains unchanged (note different residue numbering in both works, offset by 19 residues). We systematically employ a combination of information theory (AMINO)^25^ and statistical mechanics based (SGOOP)^24^ methods to determine an optimum reaction coordinate (RC) that describes Imatinib unbinding mechanism from the WT and N387 Abl. The RC is optimized by iterating between rounds of SGOOP and metadynamics simulations and then used in independent rounds of infrequent metadynamics^23^ to obtain dissociation kinetics.

We observe that Imatinib dissociates from WT and N387S Abl through two distinct pathways. The predominant Imatinib dissociation pathway from WT Abl is through the kinase hinge region, while against N387S Abl Imatinib dissociates via the *α*C helix region. Subsequently, comparing the Imatinib unbinding from the N387S and WT infrequent meta-dynamics trajectories, we observe a diminished propensity of the mutated serine at N387 to form an H-bond with the backbone carbonyl of A399 which is one residue upstream from D400 of the DFG motif. In comparison, in the WT Abl aforementioned H-bonding interaction is conserved. Furthermore, the reduced H-bonding interaction between N387S and the DFG motif gets manifested in increased flexibility of the DFG motif. We hypothesize that the increased DFG motif flexibility as observed in N387S Abl results impairs the interaction of K290 and the DFG backbone, facilitating an outward *α*C helix motion, thus allowing for Imatinib release via the *α*C helix pathway. In comparison, in WT Abl Imatinib unbinding requires a drastic conformational change involving disruption of the H-bonding interaction of N341 from the hinge and Y272 from the P-loop, creating a pathway for Imatinib to release via the hinge pathway. We reason that comparatively larger conformational change in the Abl resulting in Imatinib release from WT as compared to N387S Abl assists in faster k_*off*_ for N387S as compared to WT Abl. This work thus represents the potential to perform such investigations in the future for diverse systems using all-atom simulations that can access pharmacologically relevant timescales with minimum human intervention and prior bias.

## ACKNOWLEDGMENTS

This work was supported by the National Science Foundation, Grant No. CHE-2044165. ZS was also supported by University of Maryland COMBINE program NSF award DGE-1632976. This work used XSEDE Bridges through allocation TG-CHE180053, which is supported by National Science Foundation grant number ACI-1548562. We also thank UMD’s Deepthought2 and MARCC’s Bluecrab HPC clusters for computing resources. We would also like to thank Pavan Ravindra and Yihang Wang for discussions, and Michael Strobel and Shashank Pant for proofreading the manuscript.

## Methods

### Molecular Dynamics

Molecular dynamics simulations were initiated from the crystal structure of Abl kinase bound to Imatinib in the DFG-in state (PDB id: 1OPJ). Protonation states of titratable residues were assigned on the basis of pKa calculations performed using PROPKA 3.1^63,64^ at pH 7. N387S mutant structure was prepared using the side-chain mutation module of MOE.^65^ Using the solution builder module in CHARMM-GUI, ^66,67^ the Abl kinase-Imatinib crystal structure was solvated with TIP3P water molecules.^68^ Finally, Na^+^ and Cl^−^ ions were added, and the system was neutralized with the ionic concentration set to 100 mM. The final simulation box comprising of Abl kinase, Imatinib, water molecules, and ions was ~55 K atoms. All simulations were performed on GROMACS2018 patched with PLUMED version 2.4^69,70^ with a 2 fs time step. Simulations were performed utilizing CHARMM36 force field^71^ for protein, TIP3P for water molecules^68^ and, CHARMM General Force Field (CGenFF)^72^ for Imatinib. Temperature and pressure were kept at 300 K and 1 bar using the velocity rescale thermostat^73^ and Parrinello–Rahman barostat.^74^ The non-bonded interactions were calculated with a 10 Å cutoff, and long-range electrostatics were calculated using the particle-mesh Ewald (PME) method.^75^

### AMINO

The basis set of OPs used in this work was determined using AMINO.^25^ AMINO reduces a large set of OPs to a minimally-redundant subset using k-medoids clustering with the following mutual information based distance metric:

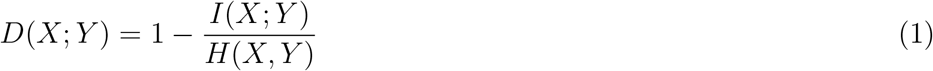

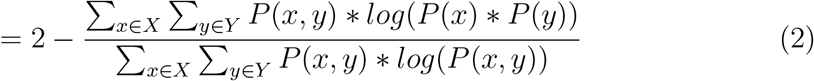

The input trajectory for AMINO was 100 ns of unbiased simulation for N387S and WT Abl bound to Imatinib with an initial set of 84 OPs corresponding to the heavy atom distance between Imatinib and the C*α* atoms of Abl. The stationary probabilities for pairs of these OPs were estimated using histograms with 50 bins and clusters of up to 20 OPs were compared to determine the optimal number of outputs. The output of AMINO was a set of 5 OPs for each system.

### SGOOP

The RCs used for enhanced sampling were linear combinations of the OPs found using SGOOP.^24,76^ This framework uses a maximum caliber model^77,78^ to construct a transition matrix along a given low-dimensional projection. The eigenspectra associated with the transition matrix is then calculated and the largest gap between consecutive eigenvalues, called the spectral gap, is found. This spectral gap represents the timescale separation between slow and fast processes. SGOOP scans different linear combinations in order to find the RC which maximizes the spectral gap and in turn the timescale separation between slow and fast processes. In this work, SGOOP was used in an iterative manner using input trajectories from the most recent simulation.

The first round of SGOOP was performed on a well-tempered metadynamics trajectory using a crude trial RC to learn a new putative RC which was a linear combination of AMINO OPs. Using the learned RC, a well-tempered metadynamics simulation was performed, to accelerate Imatinib dissociation. The resultant biased trajectory was fed back to SGOOP to learn an improved RC. The improved RC was then used to bias the next round of well-tempered metadynamics, subsequently, SGOOP was performed to further improve the RC. This process was iterated and at each step, the accelerated time and spectral gap were calculated. The iterative process was terminated when an improvement (lowering) in accelerated time was not observed (see SI Fig. 7, 9). Typically, this required 2-3 rounds of iterative SGOOP for both WT and N387S Abl systems.

**Figure 7:**
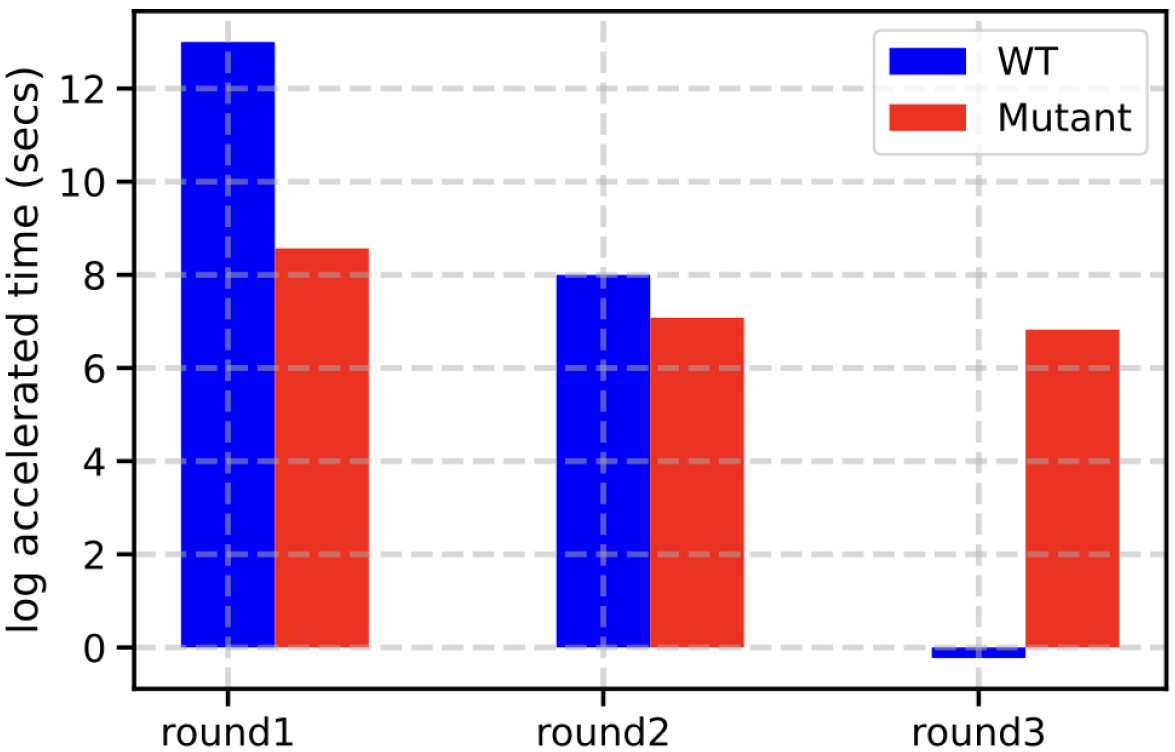
Variation in accelerated time (logarithmic scale) through rounds of preliminary metadynamics (i.e. frequent biasing as opposed to infrequent) is used to assess the quality of RC. Bar plot showing the evolution of log accelerated time in WT (blue bar) and Mutant (red bar) upon 3 stages of iterative SGOOP refinement. The reaction coordinate chosen for infrequent metadynamics run was obtained from the refinement round that corresponded to minimum accelerated time.

**Figure 8:**
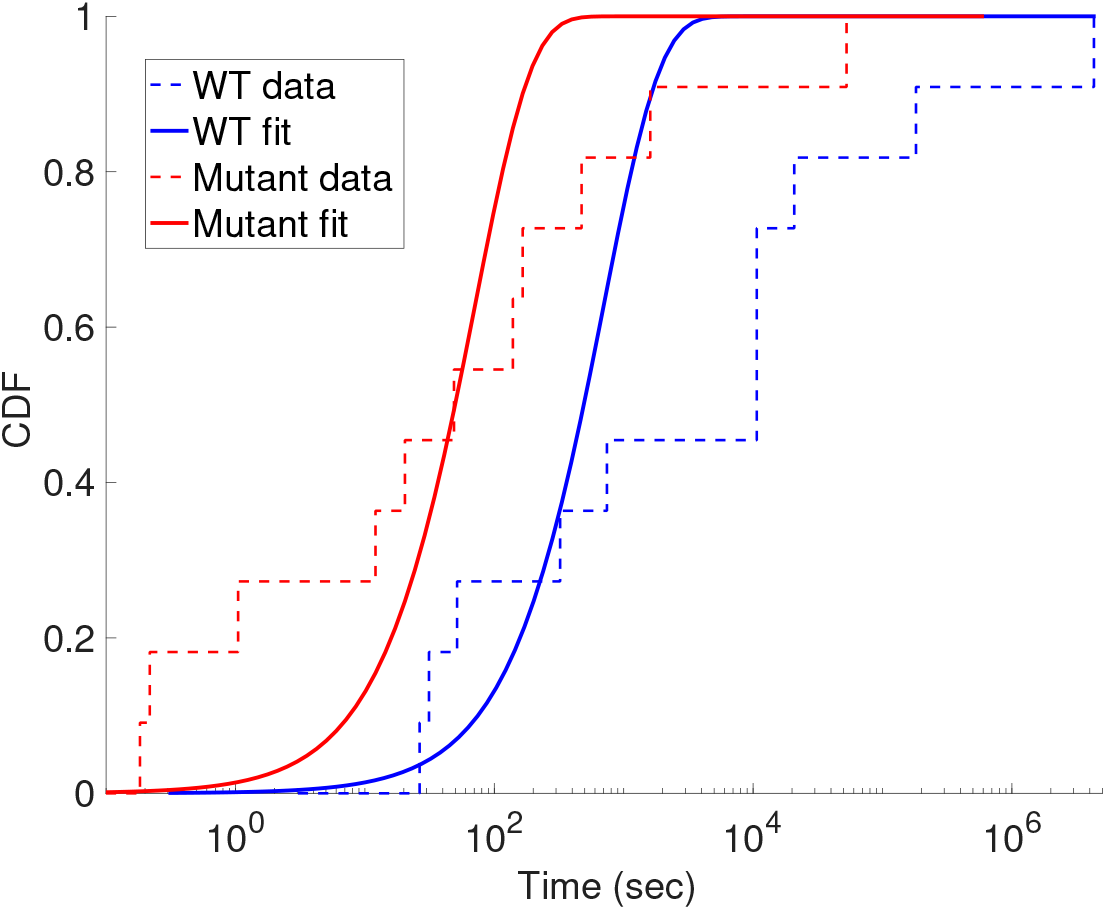
Empirical (dashed line) and fitted cumulative distribution functions (solid line) for WT (blue curves) and Mutant (red curves) corresponding to kinetics and p-values reported in main text in Table 1.

**Figure 9:**
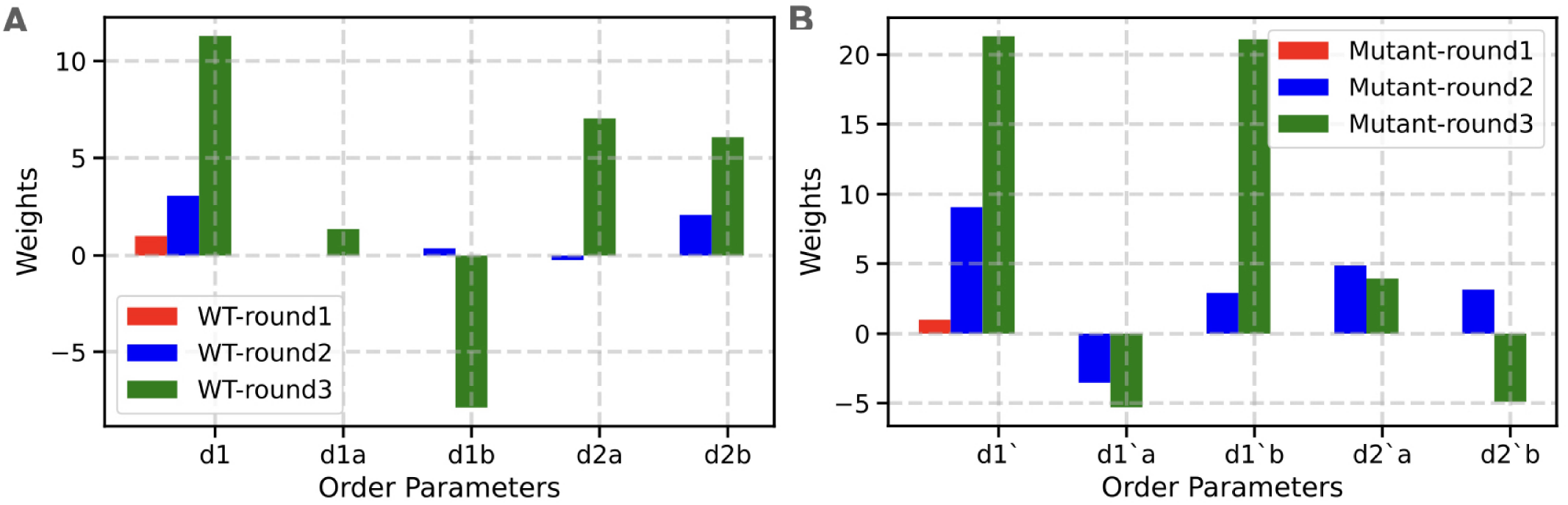
RC constructed for A) WT B) Mutant after multiple rounds of SGOOP.

### Infrequent Metadynamics

We performed 11 independent infrequent metadynamics simulations starting from the crystal structure of WT and N387S Abl. Metadynamics was performed using the PLUMED implementation of well-tempered metadynamics with a bias factor of 10, an initial hill height of kJ/mol, and bias deposited every 20 picoseconds.^69,70,79^ Biases were deposited on the RC learned from the iterative SGOOP calculation as described previously. The sigma value of the Gaussian bias kernel was estimated by calculating the standard deviation of the biased RC from the 100 ns of equilibrium MD simulation of WT and N387S Abl.

## Supplementary Information

**Table 2:** Summary of order parameters for Wild-Type and N387S simulation set. Order parameters are defined as the distance between the C*α* atom of Abl kinase and the center of mass of Imatinib atoms.

| System | OP | Abl kinase residue (Cα) | Imatinib atoms |
| --- | --- | --- | --- |
| Wild-Type | d1 | T334 | N3-N4 |
| Wild-Type | d1a | Y251 | p2a |
| Wild-Type | d1b | Y272 | p2a |
| Wild-Type | d2a | S248 | p2b |
| Wild-Type | d2b | E257 | p2b |
| N368S | d'1 | T334 | N3-N4 |
| N368S | d'1a | K293 | p2a |
| N368S | d'1b | I332 | p2a |
| N368S | d'2a | Y272 | p2b |
| N368S | d'2b | F302 | p2b |

